# Snapshot: clustering and visualizing epigenetic history during cell differentiation

**DOI:** 10.1101/291880

**Authors:** Guanjue Xiang, Belinda Giardine, Lin An, Chen Sun, Cheryl A. Keller, Elisabeth Heuston, David Bodine, Ross C Hardison, Yu Zhang

## Abstract

Epigenetic modification of chromatin plays a pivotal role in regulating gene expression during cell differentiation. The scale and complexity of epigenetic data pose significant challenges for biologists to identify the regulatory events controlling cell differentiation. Here, we present a new method, called Snapshot, that uses epigenetic data to generate a hierarchical visualization for DNA regions with epigenetic features segregating along any given cell differentiation hierarchy of interest. Different hierarchies of cell types may be used to highlight the epigenetic history specific to any particular cell lineage. We demonstrate the utility of Snapshot using data from the **VISION** project, an international project for **V**al**I**dated **S**ystematic **I**ntegrati**ON** of epigenomic data in mouse and human hematopoiesis.

**Availability and implementation: https://github.com/guanjue/snapshot**

## 1. Introduction

Multiple projects have generated thousands of epigenomic datasets, and integration of such data has become a powerful way to understand the biological meaning of the combinations of epigenetic modifications (ENCODE Project Consortium, 2012; Yue *et al.*, 2014; Bernstein *et al.*, 2010). A commonly used method for studying epigenetic patterns in multiple cell types is to first perform peak calling on individual data, and then cluster the genomic locations according to the patterns of peak presence/absence or peak intensity across all cell types (Corces *et al.*, 2016; Spencer *et al.*, 2015). These genomic locations with peaks of appropriate epigenetic features can be considered candidate cis-regulatory elements (ccREs), and their epigenetic signal pattern can reflect their role in organismal development or cell differentiation. The groups of ccREs with active epigenetic marks are often the target of transcription regulatory machinery (Huang *et al.*, 2016). Many clustering methods have been used to cluster the ccREs. The most commonly used methods are the K-means clustering method and the hierarchical clustering method (Tavazoie *et al.*, 1999; Eisen *et al.*, 1998; de Hoon *et al.*, 2004). However, these methods suffer from the assumption that the pattern of ccREs across multiple cell types are independent from each other, which is problematic because cell types are related by the process of cell differentiation. To account for the association of ccRE signals across multiple cell types, some model-based methods treat the signals of ccREs across multiple cell types as multivariate observations (Fraley and Raftery, 2002). The covariance matrix of the multivariate observations can capture the signal association across multiple cell types. More advanced methods apply either an infinite Gaussian mixture model or a Dirichlet process to further determine the number of the clusters (Rasmussen, 2000; Medvedovic *et al.*, 2004; Qin, 2006). Some other methods treat the cell types along a cell differentiation lineage as a time series and use Gaussian process mixture model to cluster “time series” data (McDowell *et al.*, 2018). However, these methods tend to group ccREs into larger clusters. As such, small clusters may be overshadowed by larger clusters. Furthermore, these methods do not consider any existing biological knowledge about the cell type relationships. Therefore, interpreting the biological meaning and visualizing the identified ccRE clusters can be difficult, especially when the number of cell types is large or when the cell types form tree-like relationships. For some of the methods, the long running time can also be an issue for large datasets.

Here, we present a different method to cluster and visualize ccREs during cell differentiation. Our method is guaranteed to identify all major clusters in the data up to a user-specified abundance threshold. The method is unsupervised and does not require the user to pre-determine the number of clusters. The method takes into account known cell-to-cell relationships in cell differentiation history. The method produces a comprehensive map of ccRE cluster patterns that lend themselves to annotation in ways that facilitate users’ ability to intuitively compare, identify, and interpret the epigenetic history during cell differentiation.

## 2. Methods

### 2.1 Clustering ccREs based on their binary indices

While unsupervised clustering methods require pre-determining the number of clusters and can miss important clusters, our method can capture all distinct and recurring clusters of ccREs. Our motivation is that each ccRE cluster corresponds to a distinct pattern of presence or absence of ccREs across the cell types examined. We first perform peak calling on all cell types using an existing peak calling method, and we call the resulting peaks “ccREs”. Next, we use the binarized presence/absence status of ccREs at each location across all cell types to create a ccRE index to represent the unique combinatorial pattern of ccREs at each genomic location (Figure 1 (a)). The number of bits in the index equals the number of cell types. The order of bits is the order of cell types derived from a user-provided cell differentiation tree. Each location with at least one ccREs across all cell types will receive an index. These indices readily classify the genomic locations into distinct ccRE clusters by assigning the ccREs with the same index to the same cluster, called an index-set (IS). Each IS contains a list of genomic locations that have the same ccRE presence/absence patterns across cell types. It is of practical utility to restrict further analyses to large ISs, i.e., the sets with many ccREs assigned to them, because the smaller ISs are often minor variations of the larger ISs. Thus we initially filter out any ISs whose size is smaller than a user-specified threshold. In the example, we choose the threshold (2500 ccREs per IS) based on the distribution of the number of ccREs per IS (Figure 1 (b)). In order to confirm that the ISs generated by grouping peak calls were supported by the initial signal (in this case for nuclease sensitivity), we analyzed the signal strength for each ccRE member of 14 ISs across five cell types. The heatmaps, generated by the deeptools package (Ramírez *et al.*, 2016), showed strong signals in the cell types for which the peak calls had indicated presence of the ccRE for ISs with numbers of ccREs above the threshold (Figure 1 (c)).

**Figure 1:**
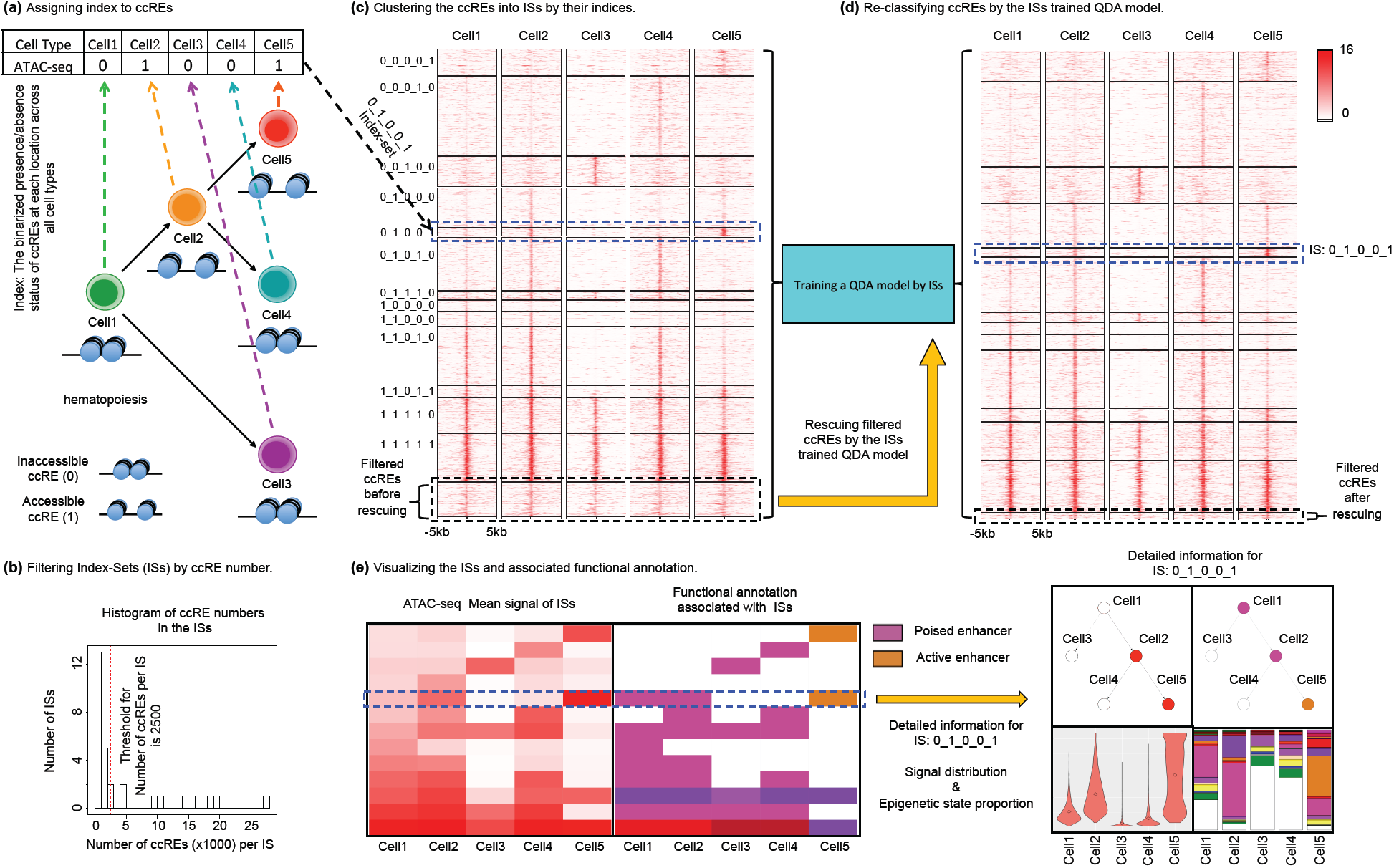
The overall workflow of snapshot. **(a)** In the 1st step, we use the binarized presence/absence status of ccREs at each location across all cell types to create a ccRE index to represent the unique combinatorial pattern of ccREs at the location. All ccREs with the same index are grouped into one cluster called an index-set (IS). **(b)** We select the threshold to filter the less prevalent ISs based on the distribution of the number of ccREs per IS (red dash line). **(c)** We visualize the ISs and their ccREs in a heatmap. The blue dash box represents one example of IS: 0_1_0_0_1. The filtered ISs and their ccREs are shown in the end of the heatmap (black dash box). **(d)** We apply the QDA model to correct ccRE’s index by borrowing information from the signals of the ccRE across all cell types. After the QDA step, some of the ccREs grouped in the rare ISs can be rescued. **(e)** Finally, we visualize the ccRE clusters in a heatmap. Each row in the heatmap is the ccRE pattern for each index-set, and each column is a cell type. The ccRE patterns are sorted by their indices in the heatmap. By our definition of the ccRE index, the ccRE patterns are separated if they have different ccRE status in a cell type; conversely they are clustered together if they have similar ccRE status in a cell type. These different plots will together highlight the epigenetic activity across cell types and the associated functional annotations and their enrichments for each index-set, which enhances the interpretation of the functional roles of each ccRE cluster during cell differentiation.

### 2.2 Rescuing ccREs by Quadratic Discriminant Analysis

The peaks detected by a peak calling method, especially for weak peaks, can be subject to errors due to data quality and the specific peak calling method used. We hypothesize that some of the ccREs in ISs whose population is below the threshold (i.e. with rare occurances) may be generated by these issues. If a peak’s signal in one cell type is associated with signals above the background at the same location in other cell types, a peak calling error at the location may be reversed by borrowing information from other cell types. We therefore developed a ccRE rescuing strategy to correct for such peak calling errors. At each genomic location with at least one ccRE in some cell type, we assume that its epigenetic signals across cell types follow a multivariate Gaussian distribution. We assume that the locations in the same IS have the same multivariate Gaussian distribution of signals, and the locations in different ISs have different signal distributions. We use all the ISs retained after filtering to train a supervised classification model called Quadratic Driscriminant Analysis (QDA) (Lachenbruch and Goldstein, 1979). We further included one “null” class in the model, which is the union of all the removed rare ISs. Then, we apply the QDA to every location in the genome that has at least one peak in some cell type to re-predict its ccRE index, which assigns the location to either an existing IS (after filtration) or to the null class. Specifically, QDA calculates the Quadratic Scores (QS) across ISs and the null class for each ccRE based on the QS function.

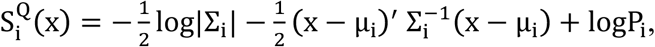

where μ_*i*_ denotes the mean vector of the *IS*_*i*_, Σ_*i*_ denotes the variance-covariance matrix of the *IS*_*i*_, *P*_*i*_ denotes the proportion of ccREs in the *IS*_*i*_, and x denotes the signal vector of each ccRE across cell types. The 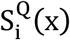 denotes the QS of *ccRE*_*i*_ for *IS*_*i*_. The QDA will assign the ccRE to the IS or null class with the highest QS. As shown in (Figure 1 (d)), a substantial number of initially filtered ccREs in initially filtered ISs are rescued by the QDA classification. For example, in a set of 158,918 ccREs predicted in five mouse hematopoietic cells, the initial filtering removed 11,954 ccREs from the ISs with membership above the threshold, and 10,048 (84%) of these were rescued by the QDA.

### 2.3 Visualizing epigenetic signal and their functional state

The ISs provide a versatile framework with which to display informative features of ccREs or other elements being analyzed. For a defining feature, such as nuclease sensitivity for ccREs, or any quantitative annotation (e.g. signal intensity for binding a co-activator, the mean signal across all elements within a set can be displayed as a unit within a heatmap (Figure 1 (e)). Each row in the heatmap shows the ccRE pattern for each IS, and each column is a cell type (Figure 1e (1)). If the annotation is categorical, such as for epigenetic states, then the most frequently occurring annotation in each IS can be displayed (Figure 1e (2)). The signal patterns are sorted by the indices of the ccREs. One common user-provided cell order places early progenitor cells at the beginning of the order and mature cells later. In this case, the ccRE patterns are more separated in the heatmap if they have different peak status in some of the “early” cell types. Conversely they are more clustered together if they have similar ccRE status in those “early” cell types. The segregation is made initially at the top of the user-provided cell hierarchy, and then it is repeated at each step down the ordered cell list. Thus the major segrating ccRE clusters for a specific cell lineage can be well separated and illustrated in the heatmap. Furthermore, a user can specify alternate lineages or cell type orders to visualize ccRE patterns in different ways. See Figure 4 and Results section for an example.

To facilitate the interpretation of ccRE clusters, our data visualization package includes four sets of plots for each IS (Figure 1 (e) (3)): (1) a cell differentiation tree colored by the intensity of epigenetic average signal in the IS in each cell type (Figure 1 (e) (3) top); (2) violinplot of the distribution of epigenetic signals in each cell type of each IS (Figure 1 (e) (3) bottom). If the user provides a whole-genome functional annotation file, our package will further generate (3) an automatically colored cell differentiation tree based on the most frequent functional annotation in the IS in each cell type (Figure 1 (e) (3) top); and (4) barplots based on the proportion of each functional annotation in the IS in each cell type (Figure 1 (e) (3) bottom). These different plots together highlight the epigenetic activity across cell types and the associated functional annotations and their enrichments for each IS. These activities and annotations facilitate the interpretation of the functional

### 2.4 Inputs and Options

We implement Snapshot as a python package with a graphical user interface. Snapshot takes the following files as input: (1) peak calling results from epigenetic data in different cell types in bed format (Kent *et al.*, 2002); (2) signal strength of the epigenetic data across the whole genome in bed format; (3) functional annotation labels in bed format (optional); (4) a list of colors for different functional annotations to be used in the heatmap (optional); and (5) a list of files containing the input file names and the corresponding content labels in the output figures. The order of the input file names in peak file name list will be used as the cell type order in the IS visualization. Snapshot uses bedtools (Quinlan, 2014) to handle most of the operations on the bed files. The user needs to provide the following parameters: (1) the minimum size (number of member elements) of an IS that should be retained (inferred to be biologically meaningful); (2) the number of rounds of QDA rescuing, which setting to 0 means clustering ccREs without peak rescuing by QDA model.

## 3. Results

### 3.1 Clustering and visualizing ccREs in hematopoietic system

We demonstrate our visualization package by analyzing the nuclease accessibility data (determined by either ATAC-seq or DNase-seq) generated and analyzed in the **VISION** project (**V**al**I**dated **S**ystematic **I**ntegrati**ON** of hematopoietic epigenomes) (Oudelaar *et al.*, 2017; Philipsen and Hardison, 2018) (Keller, Xiang, et al. manuscript in preparation). The ATAC-seq and DNase-seq data reveal genomic intervals that are accessible to nucleases, which we treat as ccREs. We first do peak calling on the nuclease accessibility data in 18 hematopoietic cell types by using the FDR adjusted p-value (*pval* _*FDR-adjusted*_<0.05) from S3norm normalization of the ATAC-seq and DNase-seq signals (Xiang, Keller, et al. manuscript in preparation). Comparing with other data normalization methods, S3norm can adjust for variation in both the sequencing depth and signal-to-noise ratio between datasets to produce comparable signals without inflating the background. We then merged the peaks in each cell into a master peak list. We used the merged master peak list (262,692 ccREs) for the downstream analysis. Next, we assigned each location with at least one ccRE an 18-digit index, where each digit corresponds to the presence (1) or absence (0) of the ATAC-seq/DNase-seq peak in each of the hematopoietic cell type (Figure 1 (a)). In theory there are 2^18^ possible combinations of the 18-digit index, but by default the heatmap tool in Snapshot plots abundant ISs containing more than 100 genomic regions (Figure 1 (c)). This threshold can filter out 11223 small ISs which contain 57508 ccREs. These ccREs will be rescued in the next step by a QDA model.

### 3.2 Snapshot re-classifies ccREs in rare ISs into abundant ISs

To rescue the filtered ccREs, we first trained a QDA model by using the information in the abundant ISs. Specifically, we used the signals across all cell type of each ccRE as the predictor and the index of ccRE’s IS as the class label to train a QDA classification model. Then we used the trained QDA model to re-classify each ccRE into the IS by its highest posterior probability (Figure 1 (d)). As such, the ccRE’s binarized presence/absence status in each cell type is decided by borrowing information of its signal across all cell types. By using the QDA model, we rescued more than 40632 (∼71%) ccREs that were filtered out in the previous run and re-classified them into larger ISs. Thus, the numbers of ccREs per IS are increased (Figure 2 (a, b)). After reclassification, many ccREs have their binarized peak status changed. For example, there are 40984 ccREs labeled as present in NK cell, among which 6139 ccREs are new to NK cell after reclassification. We used GREAT to check the biological function of these newly added NK peaks (McLean *et al.*, 2010) and obtained significant enrichment in immune system gene ontology terms (Figure 2 (c) red box). There are also 1442 peaks in the NK cell that are labeled as absent after reclassification. By GREAT, we found that these removed ccREs are in fact unassociated with immune system gene ontology terms.

**Figure 2:**
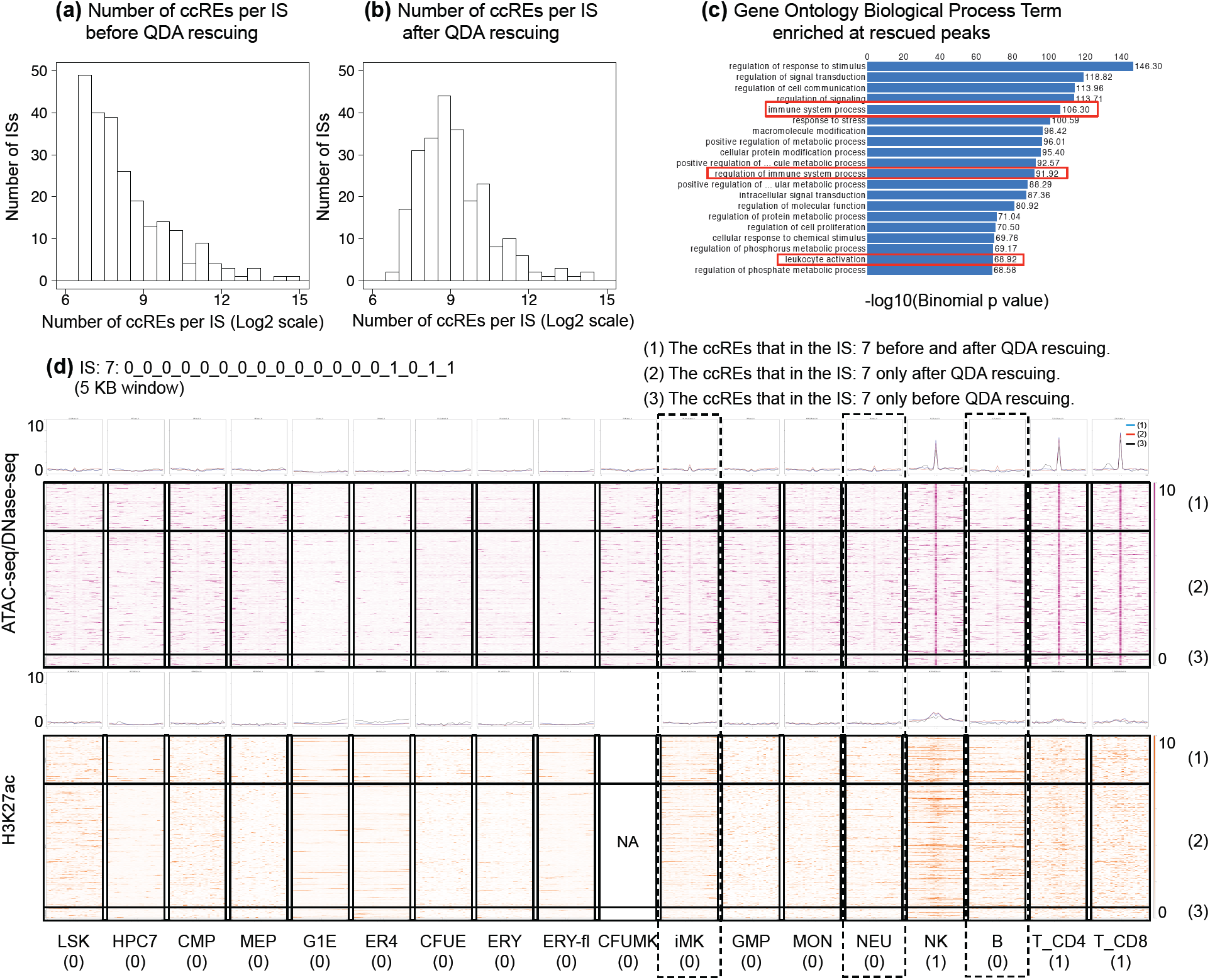
The peak rescuing by a QDA model. **(a)** The histogram of the number of ccREs per IS before QDA peak rescuing. **(b)** The histogram of the number of ccREs per IS after QDA peak rescuing. **(c)** The Gene Ontology Biological Process terms enriched in the NK cell new peaks. **(d)** The heatmap and corresponding composite plot of ATAC-seq/DNase-seq (purple) and H3K27ac ChIP-seq (orange) of IS: 7 after QDA model rescuing. There are three cluster of ccREs in the IS: 7. (1) The 1st cluster are ccREs that in the IS: 7 before and after QDA rescuing. (2) The 2nd cluster are ccREs that in the IS: 7 only after QDA rescuing. (3) The 2nd cluster are ccREs that in the IS: 7 only before QDA rescuing.

We use IS 7, which represents peaks in only NK and T cells, as an example to illustrate the newly added ccREs (Figure 2 (d)). The number of ccREs in IS 7 increased from 336 to 1067 after reclassification. From the heatmaps and the correponding composite plots (Figure 2 (d) purple heatmaps and the composite plots above them), we found that most of the newly added ccREs have weak signals in all cell types but NK and T-CD4/8 cells (Figure 2 (d) dash boxes). We further checked the H3K27ac (Active enhancer mark) ChIP-seq signal at these newly added ccREs (Figure 2 (d) orange heatmaps and the composite plots above them). These ccREs showed weak H3K27ac signals in all cell types but NK and T-CD4/8 cells as well. We thus conclude that most of the new ccREs in IS 7 are likely real. They were not originally in IS: 7 because their weaker signals in other cell types passed the threshold used for peak calling (Figure 2 (d) dash boxes).

### 3.3 Snapshot reveals clearer and more comprehensive patterns in cell differentiation system

We compared the Snapshot clustering result to the K-means clustering result (Figure 3 (b), (a)). For the K-means clustering method, we set K=20 (Figure 3 (a)). Firstly, the Snapshot clustering result is well-organized in terms of the interpretability (Figure 3 (b)). Users can easily identify the ISs and the correponding ccREs accessibility history during the cell differentiation. For example, users interested in the ccREs accessible in both common myeloid progenitors (CMP) and megakaryocytic erythroid progenitors (MEP), they can identify that one group of ccREs are accessible in erythroid (ERY) differentiation lineage (red box in Figure 3 (b)) and another group of ccREs are accessible in megakaryocytic (MK) differentiation lineage (orange box in Figure 3 (b)). It agrees with the cell differentiation tree of these cell types (Figure 3 (c)). Secondly, cell type-specific ISs can also be clearly identified, such as the IS that is only accessible in T-CD8 cell. In a sharp contrast, the K-means clustering result mainly revealed large clusters, i.e., many distinct but less prevalent clusters of ccREs are often grouped together, whereas only the distinct but common clusters of ccREs may be identified.

**Figure 3:**
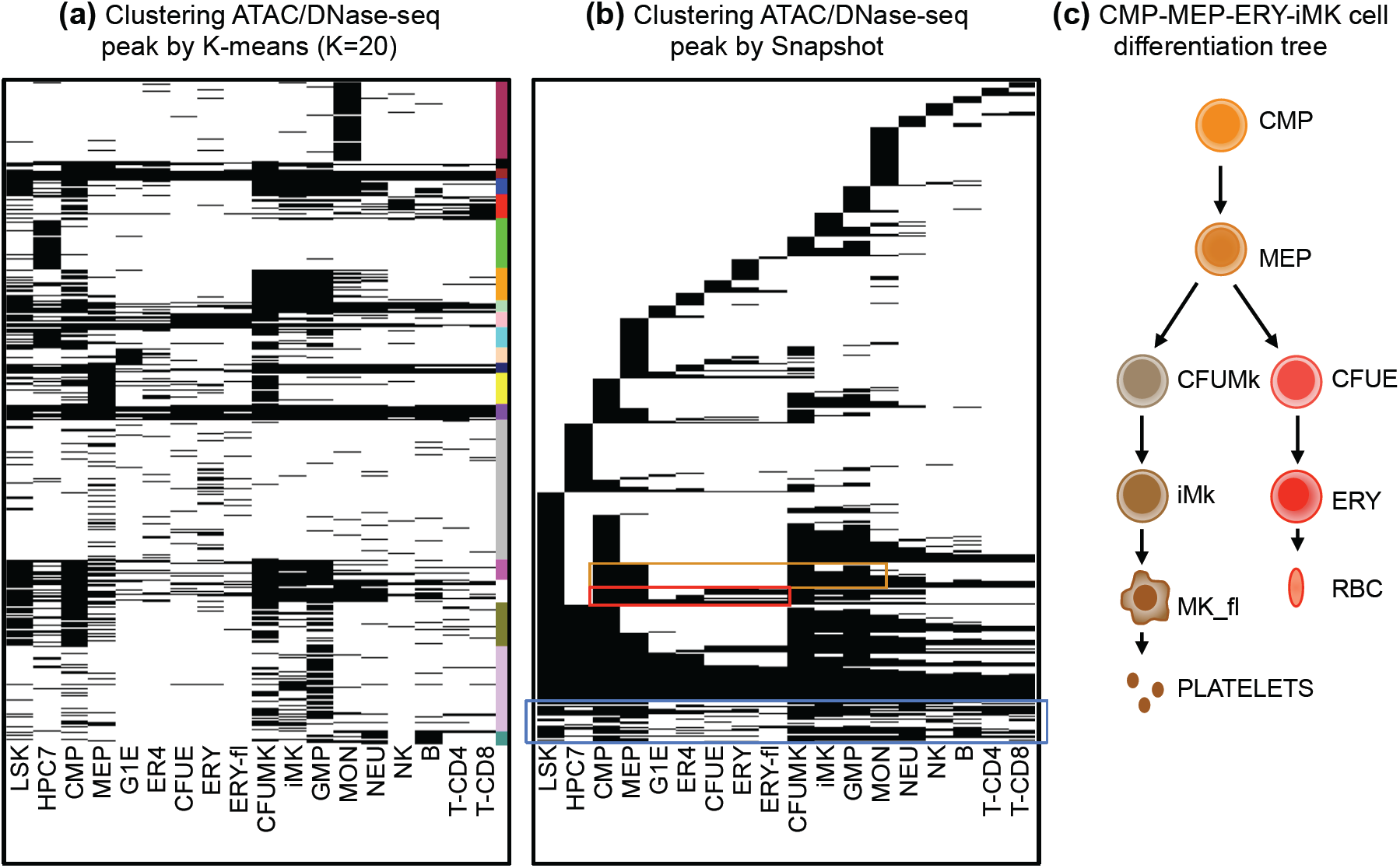
The clustering results of ccREs. **(a)** The ccREs are clustered by the K-means method. The K equals to 20. Each row represents the binarized presence/absence status of a ccRE across all cell types. The black represents the ccRE presents in the cell type and the white represents the ccRE absents in the cell type. The colorbar represents the K-means clusters. **(b)** The ccREs are clustered into ISs after QDA rescuing by the Snapshot. The ccREs in the blue box are the ccREs that are not rescued by the QDA step. **(c)** The cell differentiation tree relavant to CMP-MEP-iMK-ERY cell types.

**Figure 4:**
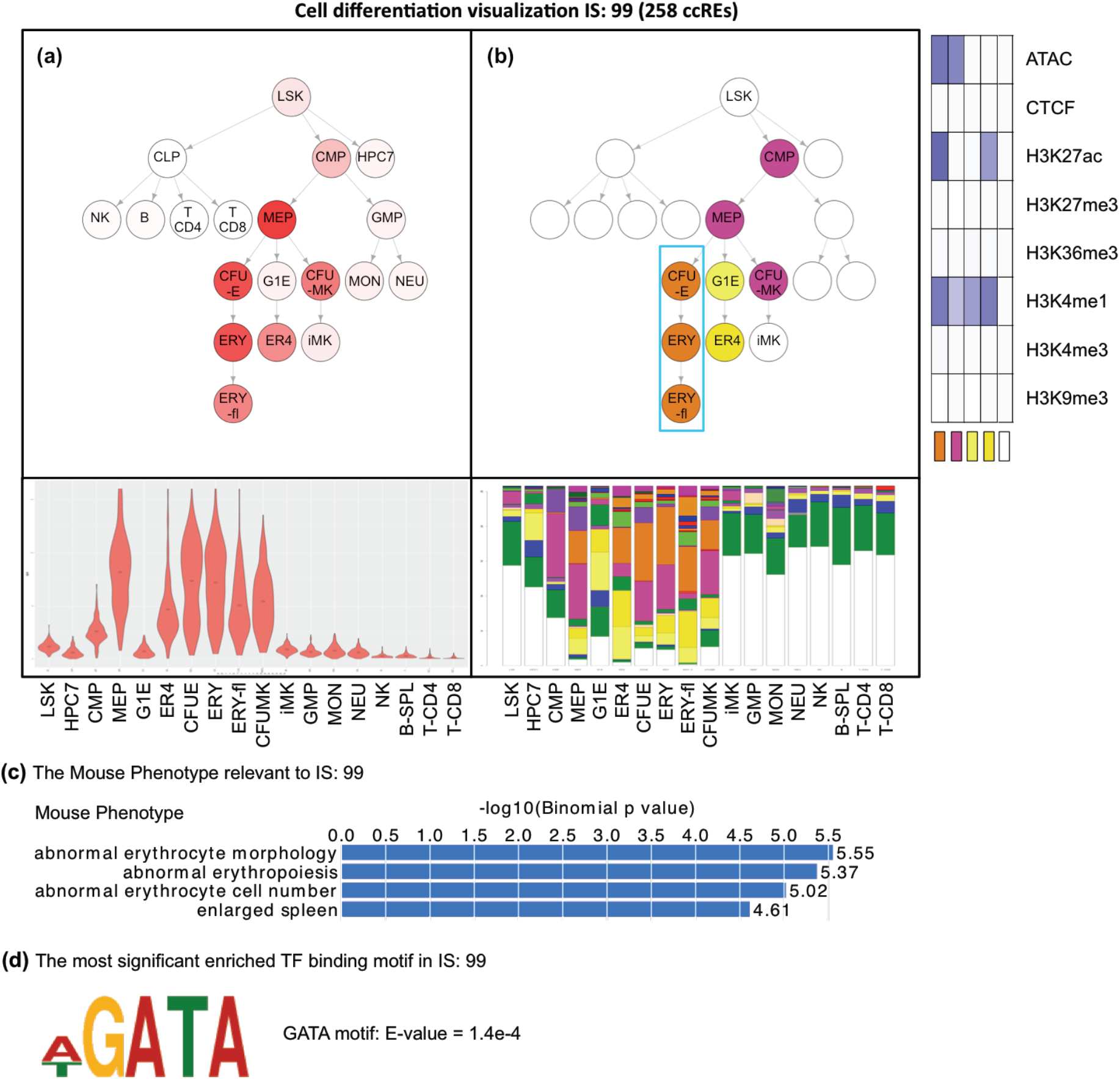
The data visualization for IS: 99 and corresponding GREAT analysis and MEME-ChIP TF binding motif analysis. **(a)** The hematopoietic cell differentiation tree colored by the average ATAC-seq/DNase-seq signal of the IS: 99 across all cell types. The violinplot represents the distribution of ATAC-seq and DNase-seq signal of ccREs in the IS: 99. **(b)** The same cell differentiation tree colored by the most frequent functional annotation in the IS: 99 across all cell types. The right heatmap represents the two most frequence functional annotations in erythroblasts lineage. The first column represents the epigenetic composition of the active enhancer state (orange). The second column represents the epigenetic composition of the poised enhancer state (purple). The barplot represents the proportion of each functional annotation in the IS: 99. **(c)** The IS: 99 significantly relevent Mouse Phenotype terms are related to erythroblasts lineage. **(d)** The most significantly enriched TF binding motif ccREs in IS: 99 from DREME analysis.

### 3.4 Snapshot helps to identify ccREs relevant to Erythroid differentiation lineage

From the four sets of plots for each IS, specific ccRE clusters (ISs) displaying ATAC-seq and DNase-seq histories of interest to the user can be discovered. For example, IS 99, an index-set containing 258 genomic regions, suggests that its ccREs may be involved in erythroid gene activation by its ATAC-seq/DNase-seq signal and functional annotation features. Specifically, the accessibility (ATAC-seq/DNase-seq signal) of this IS gradually increased from the progenitor cells to the erythroblasts (Figure 4 (a)), and its most frequent functional state became active enhancer state (orange color) upon entering the erythroid differentiation lineage (Figure 4 (b)). The functional state annotation were generated by the IDEAS 2D genome segmentation method (Zhang *et al.*, 2016) using the epigenomic data in the VISION project. These observations suggested the hypothesis that these ccREs are critical for erythroid differentiation. This hypothesis predicted that functional ontology terms for gene likely to be regulated by this set of ccREs should be enriched in terms for erythropoiesis and that the ccREs would be enriched in binding site motifs for erythroid transcription factors. To test these predictions of our hypothesis, we examined the Mouse Phenotype terms of genes associated with these regions using GREAT (McLean *et al.*, 2010). The results confirmed that the ccREs in IS 99 were significantly associated with erythroid differentiation (Figure 4 (c)). Furthermore, the most significantly enriched transcription factor binding motifs (from DREME) (Bailey, 2011; Machanick and Bailey, 2011) were those for the GATA transcription factor family. In particular, two GATA factors, GATA1 and GATA2, are critically important for erythroid cell differentiation (Katsumura and Bresnick, 2017).

## 4. Discussion

We introduce a novel tool called Snapshot that automatically generates clusters of ccREs (or other elements) in a manner that readily maps onto a cellular progression, such as a differentiation series. This simple, index-based clustering easily reveals lineage-specific or stage-specific epigenetic events. Additional tools allow users to visualize distributions of defining or informative epigenetic features, which in turn helps users formulate data-driven hypotheses about their the regulatory functions associated with the ccREs. The method does not require the user to predetermine the number of clusters, and it guarantees identification of all abundant ccRE patterns. The results are more comprehensive and readily interpretable than those from conventional clustering methods. The QDA model for borrowing information from other cell types to correct errors in peak calling further helps to identify ccREs across cell types in a robust way.

The data visualization module of Snapshot enables users to associate ccRE clusters with informative functional annotations, which can be genome segmentation results from epigenetic marks, ChIP-Seq data on protein-DNA interactions, and/or sequence information such as TF binding motifs. Furthermore, the ISs identified by Snapshot can be used as initial centers for other complex clustering methods.

While we have described Snapshot in terms of its utility for analyzing ccREs across cell differentiation, the tool can also be applied to study any progression of cell types, such as in response to hormones or signaling factors or along a developmental series. In summary, Snapshot can facilitate the discovery and interpretation of ccREs that are critical for lineage specific cell development.

## ACKNOWLEDGEMENTS

The work is supported by NIH grants GM121613 and DK106766.

## References

Bailey, T.L. (2011) DREME: Motif discovery in transcription factor ChIP-seq data. Bioinformatics.

Bernstein, B.E. et al. (2010) The NIH roadmap epigenomics mapping consortium. Nat. Biotechnol.

Corces, M.R. et al. (2016) Lineage-specific and single-cell chromatin accessibility charts human hematopoiesis and leukemia evolution. Nat. Genet.

Eisen, M.B. et al. (1998) Cluster analysis and display of genome-wide expression patterns. Proc. Natl. Acad. Sci. U. S. A.

ENCODE Project Consortium (2012) An integrated encyclopedia of DNA elements in the human genome. Nature.

Fraley, C. and Raftery, A.E. (2002) Model-based clustering, discriminant analysis, and density estimation. J. Am. Stat. Assoc.

de Hoon, M.J.L. et al. (2004) Open source clustering software. Bioinformatics.

Huang, J. et al. (2016) Dynamic Control of Enhancer Repertoires Drives Lineage and Stage-Specific Transcription during Hematopoiesis. Dev. Cell.

Katsumura, K.R. and Bresnick, E.H. (2017) The GATA factor revolution in hematology. Blood.

Kent, W.J. et al. (2002) The Human Genome Browser at UCSC. Genome Res.

Lachenbruch, P.A. and Goldstein, M. (1979) Discriminant Analysis. Biometrics.

Machanick, P. and Bailey, T.L. (2011) MEME-ChIP: Motif analysis of large DNA datasets. Bioinformatics.

McDowell, I.C. et al. (2018) Clustering gene expression time series data using an infinite Gaussian process mixture model. PLoS Comput. Biol.

McLean, C.Y. et al. (2010) GREAT improves functional interpretation of cis-regulatory regions. Nat. Biotechnol.

Medvedovic, M. et al. (2004) Bayesian mixture model based clustering of replicated microarray data. Bioinformatics.

Oudelaar, A.M. et al. (2017) Between form and function: The complexity of genome folding. Hum. Mol. Genet.

Philipsen, S. and Hardison, R.C. (2018) Evolution of hemoglobin loci and their regulatory elements. Blood Cells, Mol. Dis.

Qin, Z.S. (2006) Clustering microarray gene expression data using weighted Chinese restaurant process. Bioinformatics.

Quinlan, A.R. (2014) BEDTools: The Swiss-Army tool for genome feature analysis. Curr. Protoc. Bioinforma.

Ramírez, F. et al. (2016) deepTools2: a next generation web server for deep-sequencing data analysis. Nucleic Acids Res.

Rasmussen, C.E. (2000) The Infinite Gaussian Mixture Model. Adv. Neural Inf. Process. Syst. 12.

Spencer, D.H. et al. (2015) Epigenomic analysis of the HOX gene loci reveals mechanisms that may control canonical expression patterns in AML and normal hematopoietic cells. Leukemia.

Tavazoie, S. et al. (1999) Systematic determination of genetic network architecture. Nat. Genet.

Yue, F. et al. (2014) A comparative encyclopedia of DNA elements in the mouse genome. Nature.

Zhang, Y. et al. (2016) Jointly characterizing epigenetic dynamics across multiple human cell types. Nucleic Acids Res.

